# Predicting Macroscopic Axon Topology from Microscopic Kinematics: An Interactive Tracking and Random Walk Pipeline for Substrate-Dependent Cortical Neurospheres

**DOI:** 10.64898/2026.07.30.741748

**Authors:** Chunghwan Kim, Myungbo Kim, Hanlin Cao, Tsung-yeh Hsieh, Yongjie Jessica Zhang, Tzahi Cohen-Karni, Victoria A. Webster-Wood

## Abstract

The cortical neuron is a fundamental building block of the mammalian brain, and the morphology of its axonal projections is central to how functional circuits assemble. The trajectory along which an axon grows is a key determinant of connectivity, yet the kinematics of cortical axon outgrowth remain poorly quantified. Characterizing these dynamics is most tractable *in vitro*, where axonal growth can be measured directly and under controlled, reproducible conditions. Even in culture, however, this remains challenging because cortical neurons require dense plating for viability, and their soma is motile, so growth behavior is highly sensitive to local density and population context, complicating reproducible measurement of intrinsic dynamics. To overcome these limitations, we used size-controlled cortical neurospheres, which provide a fixed spatial origin and a reproducible environment, together with a custom semi-automated tracking pipeline to quantify single-axon kinematics across two functionalized substrates and two developmental phases. This approach revealed a substrate-dependent divergence in outgrowth: during the later developmental phase, axons on poly-D-lysine with laminin (PDL-LA) substrate grew faster than those on PDL, with a mean step size of 0.436 versus 0.339 ***µ***m /min. Decomposing trajectories into Katz dynamic states, we built a generative biased random walk model that reproduces axonal behavior at both microscopic (single-axon) and macroscopic (network topology) scales. This open, reproducible framework links single-axon kinematics to network architecture, enabling the structural connectivity of neurosphere-based circuits *in vitro* to be predicted from measurable growth dynamics, a necessary foundation for future studies linking circuit structure to emergent function.

## 1 Introduction

The computational capacity of the mammalian brain is fundamentally governed by its intricate morphological architecture [1–3]. Unlike typical somatic cells, neurons can contain more than 99% of their cytoplasmic volume within highly arborized dendritic and axonal projections (collectively, neurite structure) [4]. The neuronal axons can span remarkable distances within the brain, ranging from hundreds of micrometers between adjacent cortical areas to centimeters across hemispheres or along long-range projection pathways such as the corticospinal tract, with the exact length scale depending on species and circuit [5–8]. In addition, this extensive, yet highly distinct, neurite structure also allows individual neurons to establish the complex modular neuronal circuits that give rise to higher-order cognitive and sensorimotor functions [1–3, 9]. Before neurons can form functional synaptic networks, they must initiate outgrowth and project their neurites toward a distant target. Although the importance of this intimate structure-function relationship has long been recognized, a comprehensive understanding of how individual cells physically navigate to construct these complex networks remains elusive.

Complex navigation is driven by neuronal growth cones at the tip of the axon, which play a critical role in guiding the extending neurites to their appropriate targets during neural development [10–13]. To understand this physical navigation, the complex dynamics of neurite growth, including elongation, retraction, lateral sliding, and continuous changes in trajectory angle, have traditionally been characterized by the tracking of growth cones over time [14–17]. However, these metrics exhibit substantial biological variability and even span a significant range. For example, overall growth cone movement rates range from 0.04 µm*/*min for *Aplysia* neurons [18] through

0.34 µm*/*min for chick sensory ganglia neurons [14, 19] to 0.47 µm*/*min for frog neurons from Xenopus embryos [20]. Furthermore, it has been shown that these overall growth rates can be influenced not only by the intrinsic identity of the cell, including host species and specific neuronal subtype, but also by neuron age [21], by substrate characteristics (e.g., substrate types [22] and substrate topography [16]), and by extrinsic microenvironment factors such as electrical fields [23] and chemoattractant gradients [24]. Consequently, strictly controlled, cell-type-specific experiments are required on defined functionalized substrates to isolate true intrinsic growth from ambient noise.

The quantitative kinematics of axonal growth *in vitro* have been well established in peripheral sensory neurons (e.g., Dorsal Root Ganglia) [14, 19, 25, 26] and invertebrate neurons [18] that exhibit rapid, highly directional, and robust outgrowth. In contrast, the exact growth dynamics of mammalian cortical neurons remains relatively underexplored. This gap persists primarily because mammalian central nervous system (CNS) neurons present two distinct biological hurdles that complicate quantitative analysis of mammalian CNS dynamics. First, the density-dependent viability of cortical neurons inherently complicates kinetic tracking. *In vitro* survival and extensive neuronal development require dense plating (e.g. 8.8 × 10^5^ cells/cm^2^), restricting theoretical intercellular spacing to roughly 10.7 µm [27]. Given that cortical axons extend hundreds of microns to millimeters [28, 29], this proximity causes rapid spatial overlap, making long-term isolation of individual growth dynamics nearly impossible in standard cultures. Second, cortical neurons undergo continuous somatic migration, which constantly changes the base reference coordinates of the extending axon [30]. Standard fixed-coordinate tracking algorithms fail to account for this dynamic origin, frequently misinterpreting somatic drift as true axonal elongation and introducing artifacts into kinetic datasets. Although some methodologies attempt to isolate somatic movement by physically compartmentalizing cultures within polydimethyliloxane (PDMS) microfluidic devices, imposing such artificial geometric constraints or topography introduces a new set of artifacts [31–33]. Spatial features exert contact guidance (thigmotaxis) on the mechanosensitive growth cone [34], artificially enforcing linear [35], wall-directed trajectories [33]. Consequently, these physical boundaries restrict the number of extending axons and structurally suppress intrinsic lateral exploration and collateral branching [35, 36].

Overcoming these limitations to enable clean, artifact-free measurement of cortical axon growth is not merely a technical concern; it is a prerequisite for a broader goal that has become increasingly central to neuroscience and neural engineering: the ability to rationally design and build neuronal circuits with defined architecture [37]. Engineered *in vitro* neural circuits provide a bottom-up route to test how specific structural motifs give rise to network-level dynamics and computation, an approach that has recently gained momentum through demonstrations that cultured neuronal networks can learn, adapt, and perform goal-directed tasks [38–40]. Studying these circuits *in vitro* rather than *in vivo* offers decisive advantages: cellular composition, connectivity, and extracellular environment can be precisely specified and manipulated; growth and activity can be continuously monitored at single-axon resolution using optical and electrophysiological methods that are difficult to apply in the intact brain; and systemic confounds are eliminated, allowing reproducible controlled interrogation of structure-function relationships [41, 42]. Realizing this potential, however, depends on the ability to predict how a given circuit design will develop because the trajectories along which axons grow determine which neurons ultimately connect and thus the structural scaffold on which function emerges. A quantitative and predictive understanding of axonal growth dynamics is, therefore, a prerequisite for designing engineered neural circuits with intended connectivity.

In this study, we address these challenges using size-controlled mouse cortical neurospheres as a tractable model to quantify axonal growth dynamics. The neurosphere provides a fixed spatial origin and a reproducible local density, directly overcoming the motile-soma and density-dependence limitations that complicate dynamic analysis in dissociated cortical cultures. To extract single-axon kinematics with high fidelity, we have developed a semi-automated interactive tracking pipeline that combines image enhancement, drift compensation, and operator-supervised quality control, enabling accurate trajectory extraction even under low contrast and in the presence of structural branching. Using this pipeline, we have characterized neurite kinematics on two widely used functionalized substrates (Poly-D-Lysine, PDL or Poly-D-Lysine with Laminin, PDL-LA) and across two developmental phases of cortical neuron growth *in vitro*. Building on these measurements, we have constructed a generative biased random walk model that encodes the measured axonal kinematics within a two-tiered hierarchical structure, capturing both axon-to-axon and step-to-step variability. The model was validated at two scales: against immunostained neurospheres for macroscopic axonal network topology, and against the measured single-axon kinematic distribution for microscopic dynamics. By open-sourcing the complete analysis, tracking pipeline, and generative model together with the extracted datasets, we provide a quantitative and reproducible framework that can serve as a design guideline for the rational engineering of neuronal circuits.

## 2 Methods

### Primary cell culture

Cortical neurons were obtained from primary embryonic day 18 (E18) mouse dissociated cortical tissue from BrainBits LLC (SKU: C57EDCX). For cortical neuron plating, the dissociated tissue was resuspended by titrating it five times using a sterile 1 mL micropipette with the tip cut approximately 2 mm from the original orifice to create a larger diameter. The cells were then centrifuged at 200 rpm for 1 min and re-suspended in 1 mL of fresh Neurobasal Plus medium (catalog no. A3582901, ThermoFisher) supplemented with 1× B27 Plus (catalog no. A3582801, ThermoFisher) and 1% GlutaMAX (catalog no. 35050061, ThermoFisher). The number of cells was calculated by adding 20 µL of the cell solution to 80 µL of Trypan Blue (catalog no. 15250061, ThermoFisher). 80 µL of the Trypan Blue cell solution was added to a hemocytometer to calculate the number of cells in the solution.

### Neurosphere formation

A 24-well plate of AggreWell 800 (catalog no. 34811, StemCell Technologies) was prepared by pre-treating wells with 500 µL of 1% w/v Pluronic F-127 (catalog no. P2443, Sigma-Aldrich) in deionized (DI) water and sterile filtered with a Nalgene Rapid-Flow sterile disposable filter (catalog no. 568-0010, ThermoFisher). The wells were then centrifuged at 1300 g for 5 minutes and observed under a microscope to ensure that no bubbles were trapped in any microwell. The pluronic F-127 solution was then aspirated from the well, and each well was rinsed with 2 mL of warm Neurobasal Plus medium supplemented with 1× B27 Plus and 1% GlutaMAX. After aspirating the medium during the final rinse of the well, 1 mL of the same warm medium was added to each well to be used. The prepared mouse cortical neurons E18 were then seeded in wells and carefully titrated three times to ensure a uniform distribution in the microwells. For this experiment, each microwell is estimated to contain 7,300 cells. Warm media was added to each well to achieve a final volume of 2 mL. The AggreWell plate was then incubated at 37°C with 5% CO_2_. Half media changes occurred every 3 days.

### Neurosphere transfer

24-well plates were prepared for coating with PDL using O_2_ plasma treatment at 50W for 1 min (IPC Barrel Etcher, Branson). After treatment, 1 mL of 70% ethanol was added to sterilize the dishes. Dishes with 70% ethanol were left in the biohood with the UV light on for 1 hour for further sterilization. After sterilization, remaining ethanol was pipetted out, and the wells were rinsed with 1 mL of sterile DI water three times. 50 µg*/*mL PDL solution (catalog no. A3890401, Gibco) was added to the wells and left to incubate in the biohood for at least an hour. After incubation, the PDL solution was removed with a micropipette, and the wells were washed with 1 mL of 1× Dulbecco’s phosphate-buffered saline (DPBS, catalog no. 14190250, Gibco) three times. 20 µg*/*mL mouse LA solution (catalog no. L2020, Sigma-Aldrich) was added to half of the wells and left to incubate at 37°C with 5% CO_2_ for at least an hour. After incubation, the mouse LA solution was removed with a micropipette, and the wells were washed with 1 mL of 1× phosphate-buffered saline (PBS, catalog no. 10010023, Gibco) two times and lastly washed with 1 mL of media. Media was then removed, leaving only enough media to cover the wells of the 24-well plate. Neurospheres were individually transferred on day *in vitro* (DIV) 6 onto the well plate and incubated at 37°C with 5% CO_2_ for 30 minutes before 2 mL of warm media was added to the wells. Half media changes with warm media occurred on DIV 9 as shown in Figure 1.

**Fig. 1.**
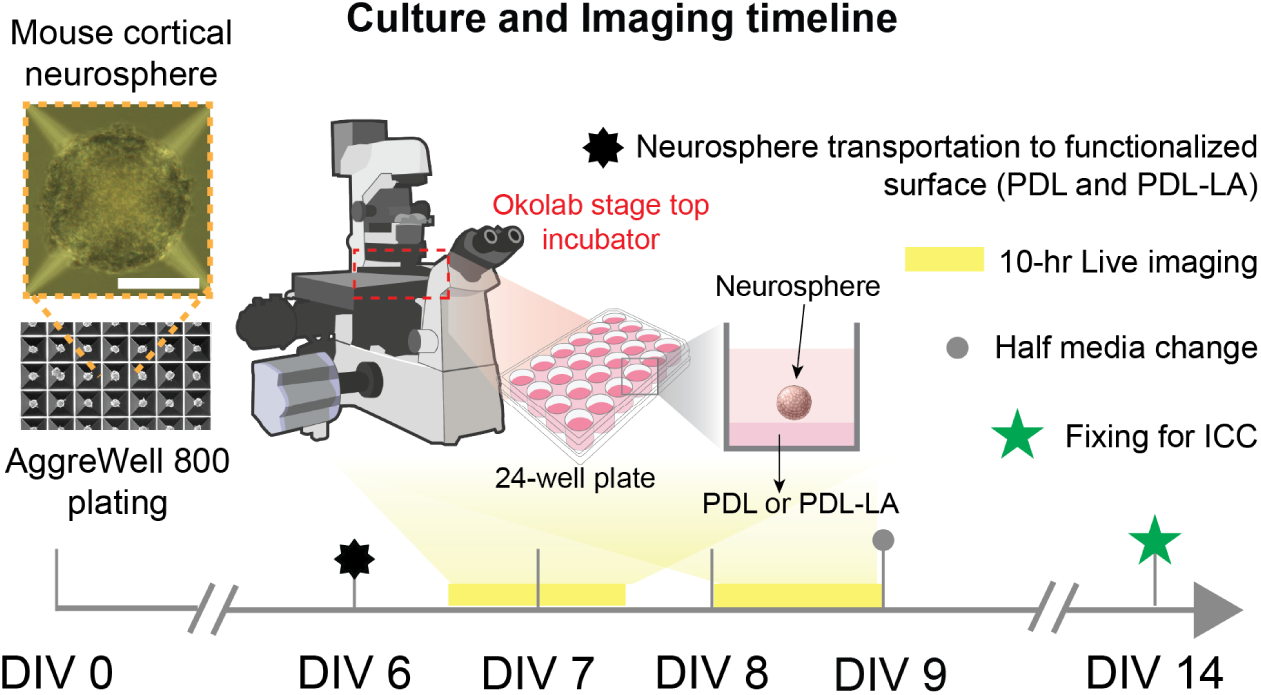
Experimental timeline for mouse cortical neurosphere culture and live-cell imaging. Mouse cortical neurospheres were initially formed using an AggreWell 800 plate. At day *in vitro* (DIV) 6, neurospheres were transferred to imaging surfaces functionalized with either poly-D-lysine (PDL) or a combination of PDL and laminin (PDL-LA). Following a 12-hour incubation period to allow for tissue adhesion and initial outgrowth, the first 10-hour live-cell imaging session (Phase 1) was conducted. A subsequent 10-hour live imaging session (Phase 2) was initiated 12 hours after Phase 1 to capture sustained axonal dynamics. Finally, cultures were maintained until DIV 14, at which point they were fixed and processed for immunocytochemistry (ICC) using a *β−*tubulin marker to visualize the established axonal networks. Scale bar represents 100 *µm*.Illustrations were obtained from NIH BioArt Source and are in the public domain: NIAID Visual & Medical Arts. (10/7/2024). Super Resolution Fluorescence Microscopy. NIAID NIH BioArt Source. bioart.niaid.nih.gov/bioart/503; NIAID Visual & Medical Arts. (10/7/2024). 24 Well Plate. NIAID NIH BioArt Source. bioart.niaid.nih.gov/bioart/4; NIAID Visual & Medical Arts. (10/7/2024). Organoid. NIAID NIH BioArt Source. bioart.niaid.nih.gov/bioart/399.

**Fig. 2.**
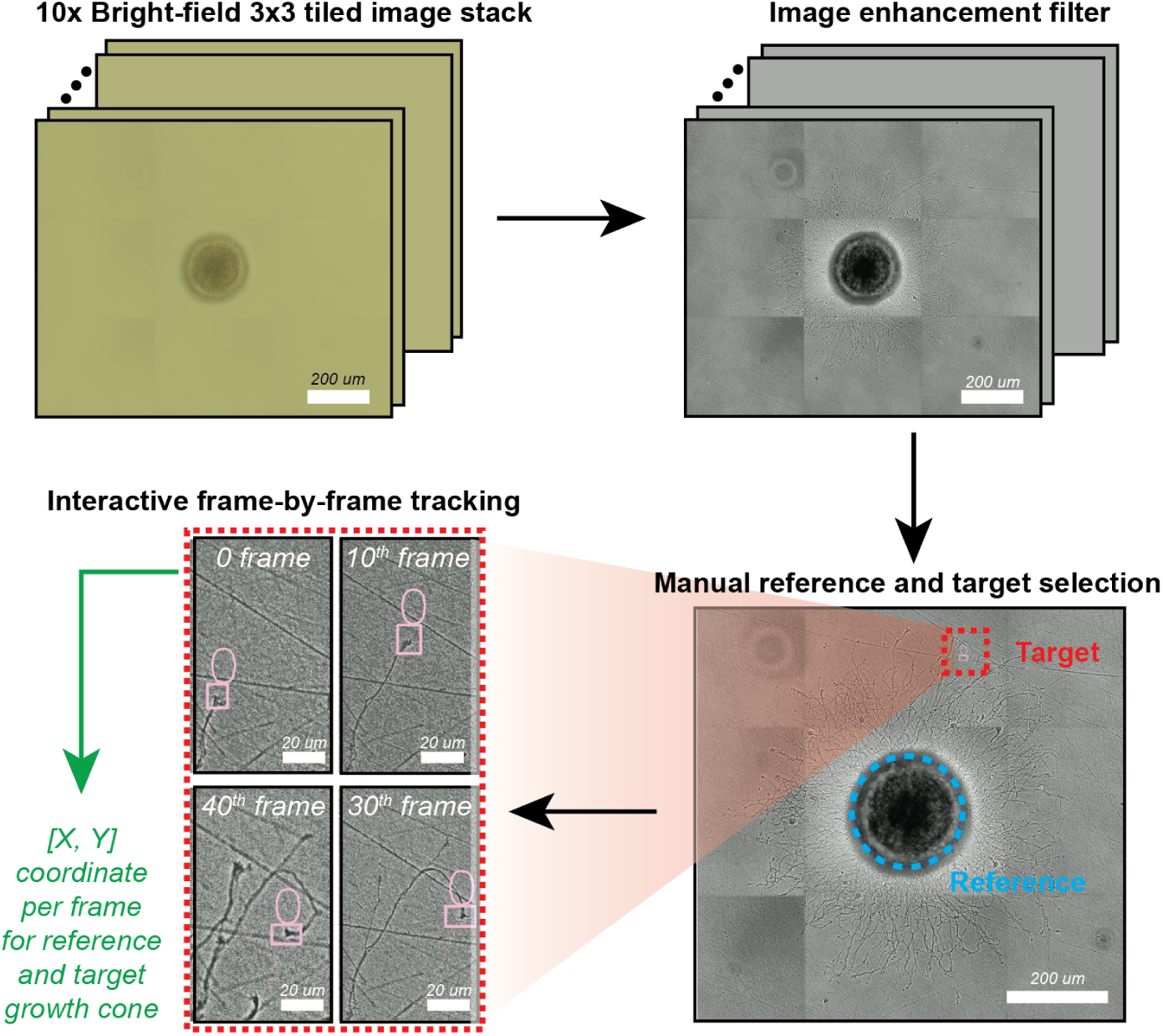
Schematic overview of the interactive neurite growth tracking algorithm. The computational workflow features a custom Python-based graphical user interface (GUI) designed for the precise extraction of raw spatial (X,Y) coordinates from longitudinal live-cell imaging data. By actively anchoring these coordinates against the center of the neurosphere (i.e., reference point), the algorithm effectively compensates for systemic image drift, thereby ensuring highly rigorous and mathematically accurate characterization of neurite kinematics.

### Live-cell imaging

To capture microscopic axonal growth, cultures were imaged on an Echo Revolution microscope in inverted bright-field mode at 20X magnification. A 3 × 3 tiled stack was acquired per well to ensure full coverage of neurite outgrowth from each cortical neurosphere transferred onto the 24-well plate. After transfer to functionalized substrates (PDL and PDL-LA), cultures were allowed to attach for 12 hours, then transferred to an Okolab stage-top incubator (H301-K-FRAME with active temperature, humidity, and *CO*_2_ controllers) mounted on the microscope for 10 hours of continuous live-cell imaging (Phase 1). The plates were subsequently returned to the standard incubator for an additional 12 hours, then returned to the stage-top incubator for a second 10-hour live-imaging session (Phase 2). The Okolab stage-top incubator maintained controlled temperature (37°C), humidity, and *CO*_2_ (5%) throughout each imaging session to preserve culture viability.

### Image preprocessing and an interactive neurite growth tracking algorithm for high-fidelity trajectory extraction

To extract single-neurite trajectories from time-lapse bright-field image stacks, we developed a semi-automated, operator-supervised tracking pipeline that combines image preprocessing, frame-by-frame object tracking with drift compensation, and interactive quality control. The pipeline was implemented in Python; the full source code and the extracted trajectory datasets used in this study are publicly available at https://github.com/CMU-BORG/CBETNeurosphereTracking-and-Analysis.git and archived at https://doi.org/10.5281/zenodo.21249734.

Each frame was preprocessed prior to tracking to enhance the visibility of fine neurite structures and suppress imaging artifacts. All preprocessing was performed in Python using the OpenCV library. Gamma correction was first applied to amplify dim cytoskeletal structures, followed by unsharp masking to sharpen structural edges. Contrast-Limited Adaptive Histogram Equalization (CLAHE) was then used to enhance local contrast across spatially heterogeneous fields, and a median filter was applied to suppress residual high-frequency noise without compromising edge integrity. On the resulting first frame, the user manually defined two categories of regions of interest through an interactive bounding-box interface: a reference structure corresponding to the central neurosphere, which served as a stable anatomical anchor for drift compensation; and one or more neurite targets to be tracked, each defined by a bounding box positioned over the axon tip of interest.

Tracking was performed frame-by-frame using the Channel and Spatial Reliability Tracker (CSRT) algorithm [43], as implemented in the OpenCV library. At each frame, the reference structure was updated first, and the centroid of its bounding box was recorded as the current drift-reference coordinate. Each neurite target was then updated independently. To compensate for global stage drift and target movement during long-duration live imaging, the position of each neurite target was expressed relative to the neurosphere centroid rather than in absolute image coordinates. The compensated displacement each frame was computed as the Euclidean distance between the neurite-target centroid and the reference centroid, yielding a drift-corrected trajectory in the neurosphere reference frame.

To handle tracker failure or new neurite branching during the imaging window, the pipeline incorporated frame-level interactive supervision. After each frame was processed, the tracking result was displayed for operator review, and three correction modes were available. First, new targets could be initialized by drawing a bounding box over a newly visible structure, allowing the pipeline to capture late-emerging neurites. Second, existing targets could be deactivated when the axon had moved out of the field of view, or the target had merged with a neighboring axon. Third, individual targets could be manually repositioned, and their trackers reinitialized at the current frame. To ensure that downstream kinematic analysis was performed on sufficiently long and well-resolved trajectories, only neurites that were continuously tracked for at least 3 hours were retained for analysis. This operator-supervised workflow maintained tracking continuity across challenging imaging conditions, including low contrast, intensity heterogeneity, and structural branching, and the resulting drift-corrected trajectories for all qualifying axons were exported in CSV format for downstream kinematic analysis.

### Kinematic metric quantification

All metrics were computed per axon and then aggregated at the population level for statistical comparison. The step size (*δ*) was defined as the Euclidean displacement between consecutive frames, and the per-axon mean step size was used to characterize each trajectory. Directionality was defined as the ratio of net displacement (start-to-end Euclidean distance) to total path length, ranging from 1 for a perfectly straight trajectory to near 0 for highly tortuous motion. The turn rate (*τ*) was computed from the frame-to-frame change in heading direction, with the per-axon global turn rate defined as the total absolute angular change normalized by trajectory duration. The mean squared displacement (MSD) was computed as a time-averaged quantity at lag intervals of 10 to 120 min in 10 min increments, and the anomalous diffusion exponent *α* was estimated from the slope of a log-log linear fit to MSD versus lag time. Population-level MSD curves were generated by averaging across all axons within each condition with 95% confidence intervals. Finally, the Katz dynamic ratios [14] quantified the proportion of each axon’s trajectory spent in three kinematic states: elongation (forward advance along the trajectory axis), retraction (backward motion along the axis), and lateral wandering (off-axis motion). At each frame, the displacement was decomposed into components parallel and perpendicular to the local trajectory axis, classified into one of the three states, and accumulated over the full trajectory. The elongation, retraction, and wandering ratios were then defined as the cumulative path length attributed to each state divided by the total path length, with the three ratios summing to one.

### Statistical analysis

All statistical analyses were performed in MATLAB with a significance level of *α* = 0.05. The distributions of step size (*δ*) and turn rate (*τ*) were tested for normality and log-normality using the Anderson-Darling test (MATLAB built-in *adtest*), applied to the raw and log-transformed data; distributions were classified as log-normal when normality was rejected for the raw data but not after log transformation. This procedure was applied at two scales: the global axon-to-axon and local step-to-step distribution, consistent with the two-tiered structure of the generative model. Since most distributions deviated from normality, between-condition comparisons were performed using the non-parametric Mann-Whitney U test (MATLAB built-in *ranksum*). Population-level distributions were visualized as violin plots generated by Gaussian kernel density estimation (MATLAB built-in *ksdensity*) bounded by the empirical data range extended by 5%, with the inter-quartile range shown as a vertical black bar, the median as a white-filled circle, and individual data points indicated with solid circle markers.

### Immunocytochemistry (ICC)

At DIV 13, neurospheres were fixed in 4% paraformaldehyde (PFA) in PBS for 15 min, washed three times in PBST (0.1% Triton X-100 in PBS), and blocked overnight in 6% BSA with 0.2% Triton X-100 in PBS. For 3D immunostaining, neurospheres were incubated for 24 h with anti-*βIII*-tubulin primary antibody (ab7751; 1:600) in blocking solution, washed three times in PBST (10 min each), and then incubated for 24 h with goat anti-mouse IgG1 cross-adsorbed Alexa Fluor 647 secondary antibody (Invitrogen, A-21240; 1:1000). Nuclei were counterstained with Hoechst 33342 (1% in blocking solution) for 30 min. Following three final PBST washes (10 min each), neurospheres were mounted in PBS containing antifade mounting medium. All processes were performed at room temperature. All images were acquired on a Nikon AXR confocal microscope.

## 3 Experimental Results

### 3.1 Neurosphere Formation and Transfer-timing-dependent Axonal Growth on Functionalized Substrates

To generate uniform mouse cortical neurospheres, high-density single-cell suspensions (2.2 × 10^6^ cells per well) were cultured using the AggreWell 800 platform. Quantitative morphological analysis revealed an initial 48-hour culture (up to DIV 2) where the size of the neurosphere remains at a plateau between DIV 1 (165.17 ± 17.51 *µ*m) and DIV 2, with no statistically significant expansion (*p-value* = 0.29) (Figure 3a). Following this initial period, the neurospheres exhibited a progressive increase in diameter, eventually reaching 213.19 ± 19.72 *µ*m. Extended observation beyond DIV 6 indicated that there was no further dimensional expansion (Figure 3a). Note that since the maximum diameter achieved up to DIV 6 remains below the critical oxygen and nutrient diffusion limit (3-400 *µ*m), the neurospheres are capable of maintaining continuous mass transport, which avoids hypoxia-induced core necrosis that limits the viability of larger organoids [44, 45]. Following this stable neurosphere formation, the induction of neurite outgrowth required transferring the neurospheres onto a 2D functionalized substrate. Individual neurospheres were transferred to individual wells in a 24-well plate containing substrates functionalized with poly-D-lysine (PDL) or PDL-laminin (PDL-LA). Because prolonged suspension can restrict baseline neuronal differentiation potential [46], we hypothesized that the specific timing of neurosphere transfer would affect the post-attachment neurite outgrowth response. To evaluate transfer-timing-dependent neurite outgrowth, neurospheres were transferred at DIV 6, DIV 7, and DIV 13 onto PDL-coated substrates. Morphological characterization of subsequent neurite outgrowth on PDL substrates revealed that extended suspension time (e.g., DIV 13) prior to transfer severely compromised the ability for robust neurite extension (Figure 3b). Consequently, to maximize macroscopic axonal outgrowth without compromising intrinsic growth behaviors, DIV 6 was established as the optimal transfer window for all downstream kinematic characterizations.

**Fig. 3.**
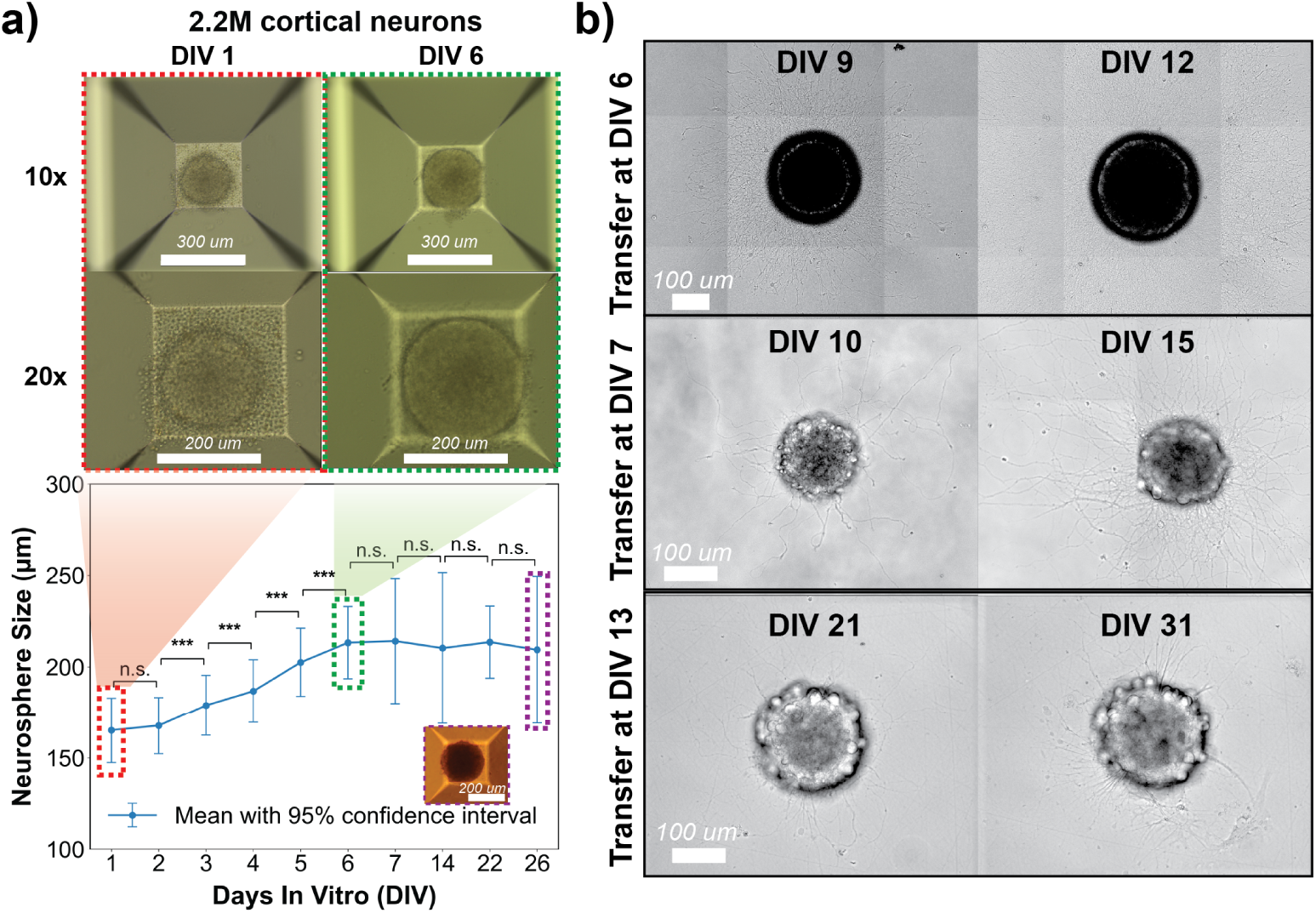
Morphological characterization of neurosphere assembly and the effect of transfer timing on axonal outgrowth. Cortical neurons were seeded at a density of 2.2*×*10^6^ cells per well in an AggreWell 800 plate to generate uniform neurospheres. (A) Representative bright-field images of neurospheres at 1 day *in vitro* (DIV 1, 10*×* and 20*×* magnification) and DIV 6 (10*×* and 20*×* magnification). The accompanying plot quantifies the progressive increase in neurosphere size from DIV 1 to DIV 26. Inset (purple dotted outline) shows a bright-field image of the neurosphere at DIV 26 acquired at 10 *×* magnification. Each condition has more than 10 samples. Statistical significance is denoted by asterisks (*** *p <* 0.005). (B) Transfer-timing-dependent axonal growth dynamics. Neurospheres were transferred to poly-D-lysine (PDL)-functionalized substrates at distinct time points: DIV 6, DIV 7, and DIV 13. Representative bright-field images obtained post-transfer demonstrate that delayed transfer onto functionalized substrates substantially restricts subsequent axonal outgrowth compared to the optimal DIV 6 transfer.

### 3.2 Spatiotemporal Kinematic Analysis of Phase- and Substrate-Dependent Axonal Outgrowth

Axonal outgrowth is a highly regulated developmental process whose complexity and kinematics are fundamentally governed by both the culture time and the biophysical properties of the underlying microenvironment [47–50]. Specifically, *in vitro* neurons transition from an initial arborizing state (*<*24 hours) to a highly directed elongating state (*>*24 hours). To effectively decouple these variables, we investigate both the influence of substrate functionalization (PDL versus PDL-LA) and the temporal evolution of axonal growth. By segmenting our live-cell tracking into two discrete developmental epochs (i.e., Phase 1: 12-22 hours post-transfer and Phase 2: 34-44 hours post-transfer), we captured a more comprehensive spatiotemporal profile of axonal growth dynamics. Although qualitative bright-field images indicate clear morphological changes between the two phases (Figure 4a), a systematic quantitative analysis is required to precisely characterize the underlying kinematics.

**Fig. 4.**
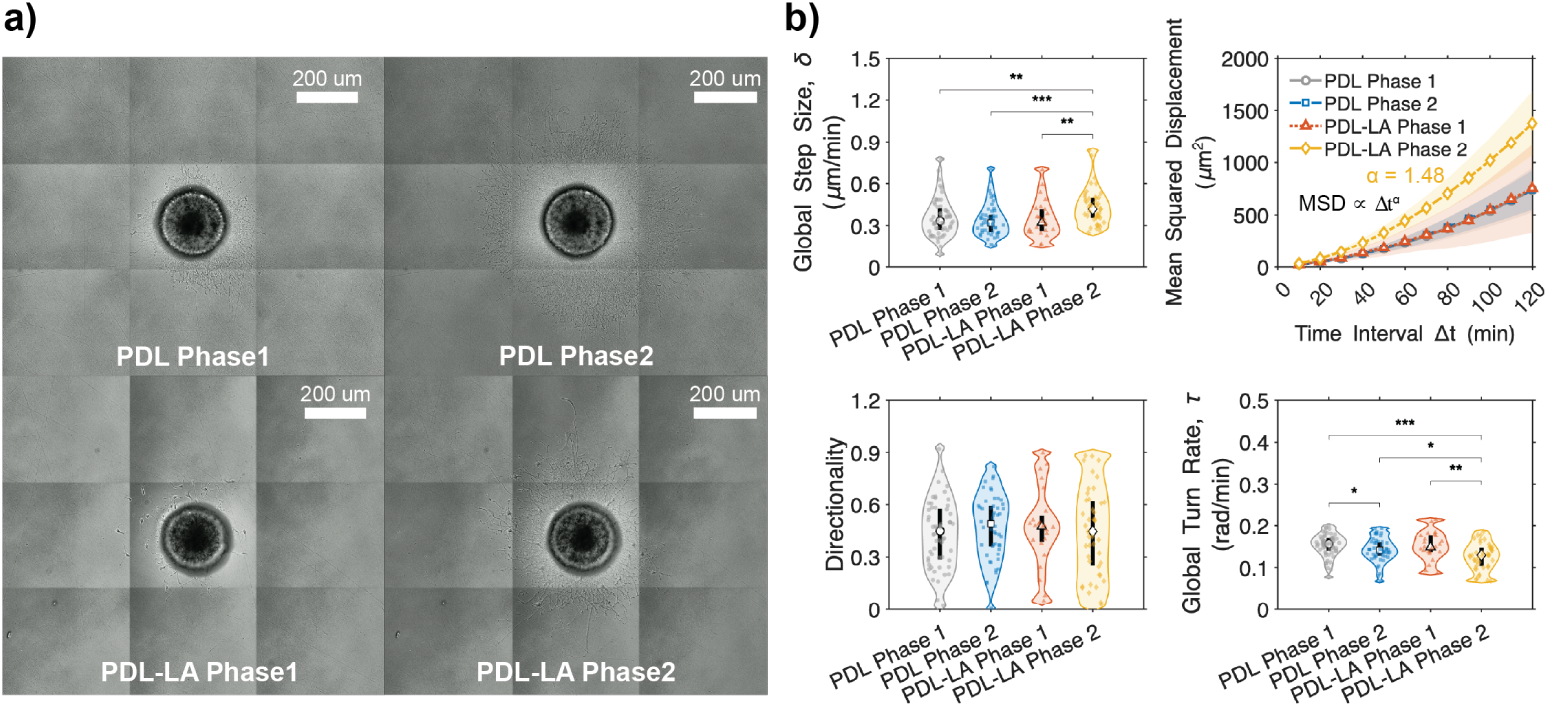
Kinematics of axonal outgrowth on PDL versus PDL-LA substrates across distinct growth phases. a) Representative bright-field images illustrating axonal extension from neurospheres cultured on poly-D-lysine (PDL) and PDL-laminin (PDL-LA) functionalized substrates during Phase 1 (12–22 hours post-transfer) and Phase 2 (34–44 hours post-transfer). b) Violin plots quantifying the macroscopic growth dynamics for each condition, with individual raw data points superimposed to display population variance. Within each violin, the central white circle denotes the median, and the thick vertical black line represents the inter-quartile range (25th to 75th percentile). Analyzed kinematic metrics include average growth step size, mean squared displacement (MSD), overall directionality, and trajectory turn rate. Statistical significance between conditions is denoted by asterisks (*: *p <* 0.05, **: *p <* 0.005 and ***: *p <* 0.001).

To achieve this, we developed a custom high-fidelity interactive tracking pipeline. This pipeline independently extracts the time-lapse coordinates of individual growth cones, mapping them directly against the neurosphere center as a fixed origin to yield precise kinematic measurements (see Methods). We first evaluated global step size (*δ*, total travel distance over total travel time), revealing that PDL-LA Phase 2 supported the fastest axonal outgrowth (0.436 ± 0.039 *µ*m*/min*), notably higher than that in Phase 1 on the same substrate (0.350 ± 0.055 *µ*m*/min*) (Figure 4b). In contrast, global speed on PDL-only substrates exhibited no significant phase-dependent acceleration (0.357 ± 0.035 *µ*m*/min* versus 0.339 ± 0.032 *µ*m*/min*) (Figure 4b). The global turn rate (*τ*), which measures the angular deviation between consecutive time steps, was minimized in PDL-LA Phase 2 (0.127 ± 0.010 *rad/min*) compared to the other conditions (0.143 − 0.153 *rad/min*) (Figure 4b). Note that the observed temporal reduction in *τ* from Phase 1 to Phase 2 captures a critical developmental shift, indicating that growth cones transition from an early exploratory state (Phase 1) into a directed axonal elongation state (Phase 2) as time in culture increases. This microscopic step-wise correlation (*δ* with highly consistent, low *τ*) motivated an investigation of macroscopic axonal growth behavior using Mean Squared Displacement (MSD). Across all conditions, the axonal growth showed a non-linear, power-law relationship with time (*MSD* ∝ Δ*t^α^*, *α* is a fitting parameter and 1 *< α <* 2), characteristic of super-diffusion [51]. PDL-LA Phase 2 showed the most aggressive super-diffusive scaling exponent (*α* = 1.48), significantly outpacing the other conditions (*α* = 1.25 −1.39) (Figure 4b). Interestingly, despite the significant increases in *δ* and *τ* in PDL-LA Phase 2, the overall directionality of the trajectory (the ratio of net displacement to the total length of the travel path) remained statistically indistinguishable under all conditions. Together, these kinematics indicate that macroscopic axonal growth acts as a super-diffusive, biased random walk, requiring the stochastic decomposition of individual trajectories to isolate the intrinsically preferred growth direction.

### 3.3 Stochastic Decomposition of Axonal Growth Trajectories

Previous studies have demonstrated that neurospheres cultured on functionalized substrates exhibit directed axonal outgrowth irrespective of the specific functionalized substrate type. To quantitatively assess this directional bias, we applied the established kinematic model developed by Katz [14]. This analytical framework mathematically decomposes complex multi-scale growth cone trajectories into three distinct stochastic vectors: directed elongation, lateral wandering, and retraction. By isolating these individual components, this stochastic decomposition enables a highly precise evaluation of the underlying macroscopic growth dynamics. In this kinematic model, an orientation of the growth cone in time step *i* establishes the primary axis of elongation (Figure 5a). Movement aligned positively with this vector defines pure elongation, while negative displacement defines retraction. Since the growth cone continuously pivots as it navigates toward step *i+1*, it simultaneously generates lateral wandering. To decouple pure elongation or retraction from this lateral wandering, the Katz model treats the trajectory between steps *i* and *i* +1 as an integration of infinitesimally small steps along the vector. Following this framework, the net longitudinal displacement (*Y*) representing either pure elongation or retraction is calculated as follows:

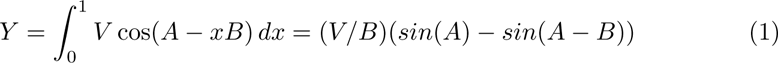

where *V* is the absolute travel distance between time steps *i* and *i* +1, *A* is the angle deviation of the travel vector relative to the primary axis of elongation, and *B* is the angle through which the growth cone pivots. The non-elongation wandering component (*W*) is defined as the residual travel distance (*W* = *V* − |*Y* |). Finally, to normalize these dynamics throughout the trajectory, individual kinematic ratios are calculated by dividing the cumulative sum of each stochastic component by the total travel distance (Σ*^V^*); the elongation ratio is based on positive *Y* increments, the retra_Σ_ction ratio on the absolute sum of negative *Y* increments, and the wandering ratio on Σ^W^ (Figure 5a)

**Fig. 5.**
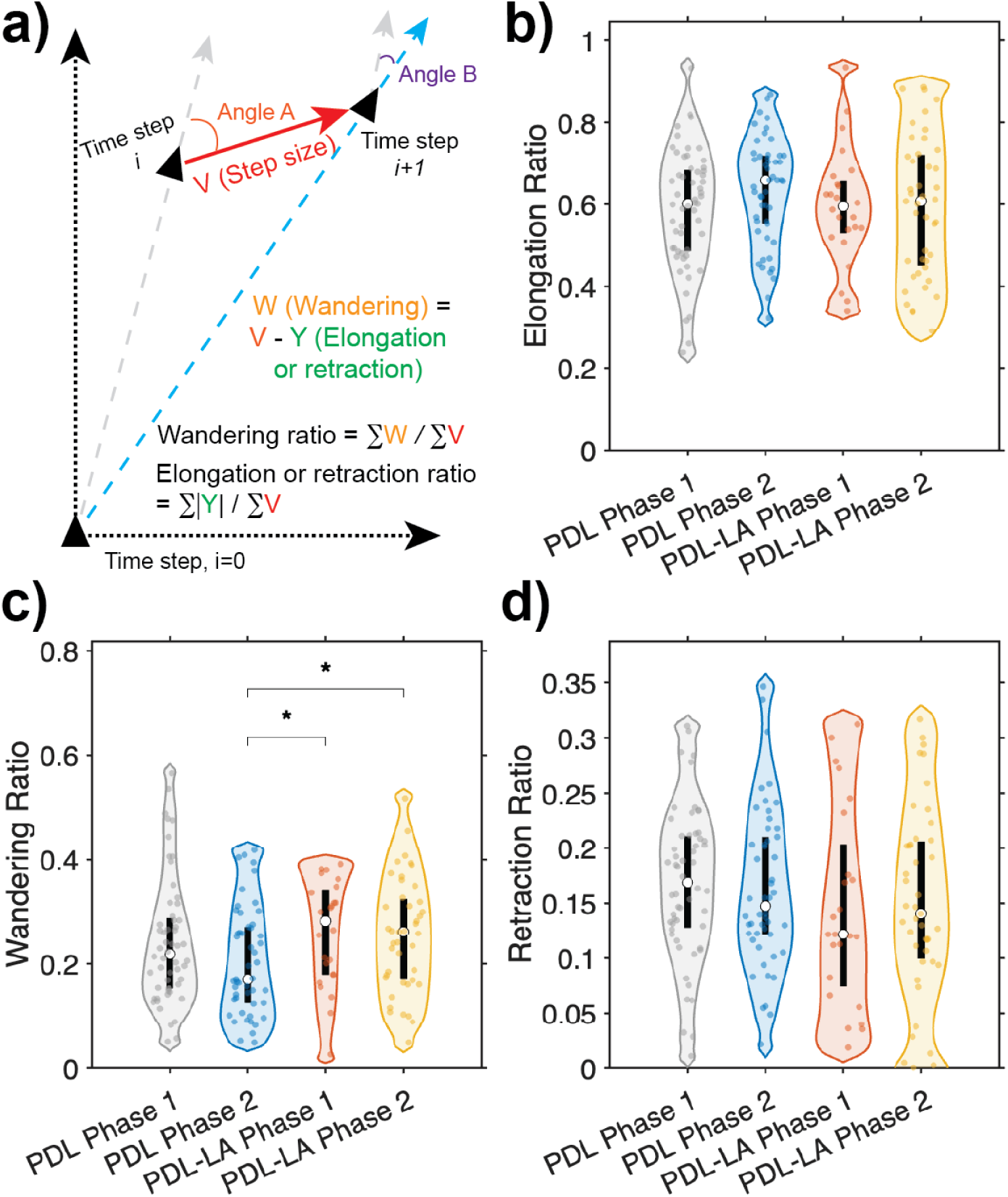
Stochastic decomposition of axonal outgrowth trajectories using the Katz kinematic model. a) Schematic diagram illustrating the decomposition of complex axonal growth trajectories into their three principal stochastic components: directed elongation, lateral wandering, and retraction. b–d) Quantitative evaluation of the Katz dynamic ratios across distinct functionalized substrates and temporal culture phases. Violin plots display the population variance with superimposed individual raw data points. Within each distribution, the central white circle denotes the sample median, and the thick vertical black line represents the inter-quartile range (25th to 75th percentile) for the b) elongation ratio, c) wandering ratio, and d) retraction ratio. Statistical significance between conditions is denoted by asterisks (*: *p <* 0.05).

Overall, the mean elongation ratio (0.59-0.63) was the highest across all phases and functionalized substrates, followed by the wandering ratio (0.21-0.26) and the retraction ratio (0.15-0.17) (Figure 5b). Notably, the mean wandering ratio for the PDL-LA substrate was consistently higher than that for the PDL substrate (Phase 1: 0.24 for PDL and 0.26 for PDL-LA, *p-value* = 0.27; Phase 2: 0.21 for PDL and 0.26 for PDL-LA, *p-value <* 0.05). The elevated wandering ratio on the PDL-LA substrate reflects a mechanistic shift from passive attachment (PDL) to active, receptor-mediated exploration (PDL-LA). By binding transmembrane integrins, laminin engages a molecular clutch that mechanically couples the extracellular matrix to the intracellular cytoskeleton [13, 52, 53]. This interaction couples F-actin retrograde flow to generate traction forces, driving the cytoskeletal remodeling and filopodial protrusions that manifest as highly exploratory axonal kinematics [13, 16, 54].

In summary, the stochastic decomposition of axonal trajectories via the Katz model confirms a highly biased dynamic regime dominated by directed elongation. This kinematic profile directly corroborates our earlier MSD analysis in Section 3.2, which characterized the macroscopic axonal outgrowth as a super-diffusive, biased random walk. By mathematically decoupling the active forward movement from lateral wandering, we have successfully isolated the intrinsic biological vectors governing this complex system. These experimentally characterized dynamics will serve as the foundational empirical parameters for the theoretical model of axonal outgrowth developed in the following section.

## 4 Theoretical Modeling

### 4.1 Generative Biased Random Walk Modeling of Cortical Neuron Axonal Growth

Early theoretical models often conceptualized growth cone motility as a pure random walk, investigating whether macroscopic elongation dynamics could emerge from stochastic processes without invoking complex intracellular mechanisms [14, 55–57]. Building upon these foundational works, we developed a generative, biased random walk model to simulate predictive axonal growth for cortical neurons *in vitro*. The initial boundary conditions of the simulation were established directly from our *in vitro* characterization. Based on nuclei quantification through Hoechst, approximately 200 cortical neurons were randomly distributed within the spatial constraints of a 220*µm* source neurosphere to initialize simulated axonal growth (Figure 6a).

**Fig. 6.**
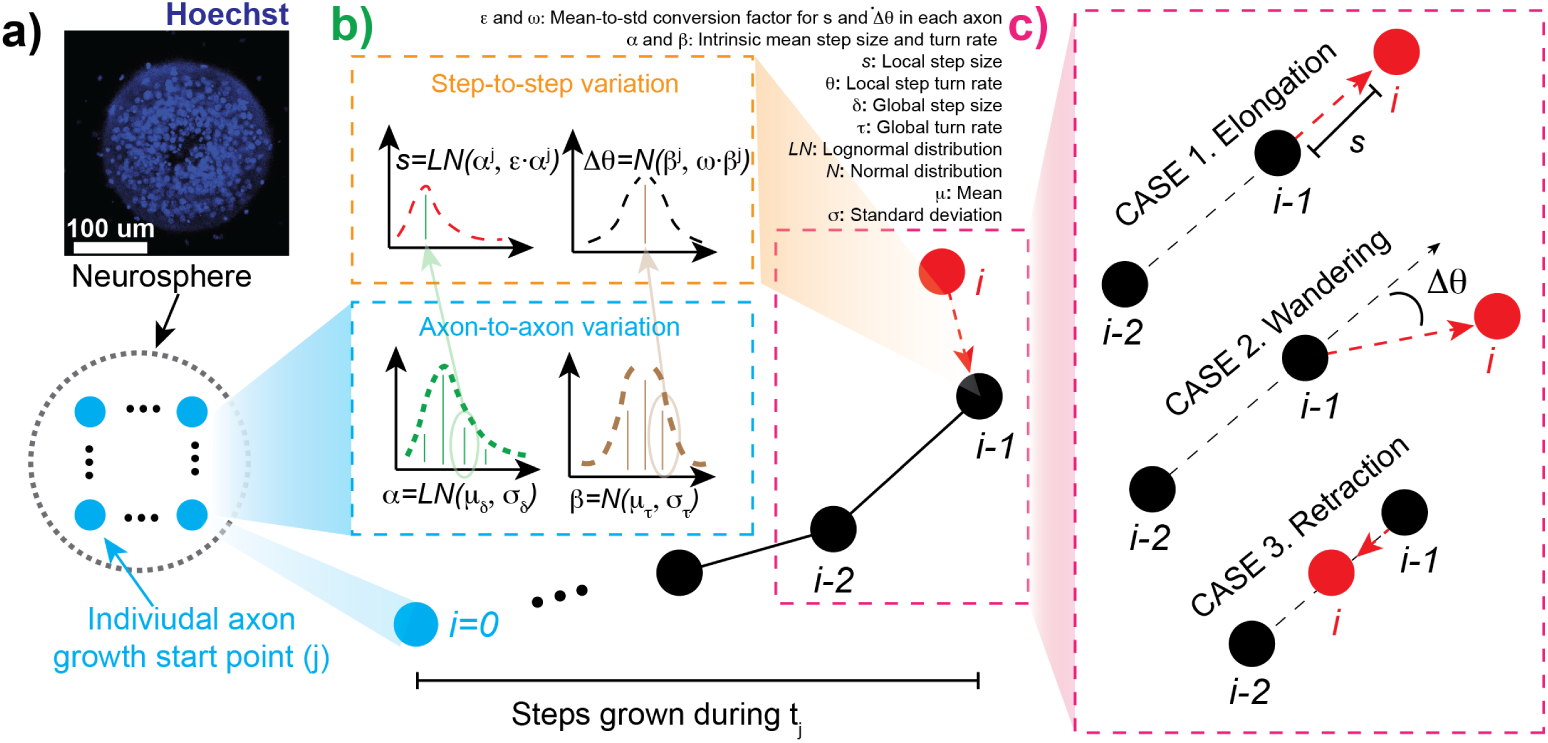
Generative stochastic modeling of axonal trajectories via a biased correlated random walk. a) Spatial initialization of the simulation. The initial coordinates for simulated axons are spatially distributed based on empirical nuclei distributions derived from Hoechst 33342 (blue) immunostaining of cultured neurospheres. b) Two-tiered hierarchy of statistical variance. The model isolates global axon-to-axon variance and local step-to-step variation. For axon-to-axon variability, each simulated axon is assigned a characteristic step size (*α*) drawn from a lognormal distribution (mean: *µ_δ_* and standard deviation: *σ_δ_*) and a characteristic turn rate (*β*) drawn from a normal distribution (mean: *µτ* and standard deviation: *στ*). For step-to-step variability, individual steps fluctuate around these baseline parameters, utilizing *ε* and *ω* as experimentally obtained mean-to-standard deviation conversion factors for step size and turn rate, respectively. c) Implementation of Katz kinematic dynamics. The simulated growth cone operates as a biased random walk that transitions between three distinct dynamic cases: CASE 1) directed elongation, CASE 2) wandering, and CASE 3) retraction. The transition probabilities governing these cases are explicitly parameterized by experimentally measured Katz ratios established in Figure 5b-d.

To produce biologically realistic axonal trajectories, the model introduces stochastic variation at two hierarchical levels: a global level, which captures variability between individual axons, and a local level, which captures step-to-step variability along each axon’s trajectory. Empirical distribution analysis of the experimental data (4b) revealed well-defined and reproducible profiles on each scale, both macroscopic (axon-to-axon) and microscopic (step-to-step) scales. Specifically, both the global step size (*δ*), defined as the average step size of each individual axon, and the local step size (*s*), defined as the step-to-step values within that axon, follow log-normal distributions (*p-value >* 0.1). In contrast, both the global turn rate (*τ*) and the individual step turn rates (Δ*θ*) follow normal distributions (*p-value >* 0.1). To simulate axon-to-axon variation, each virtual axon is assigned a unique, intrinsic mean step size (*α*) and mean turn rate (*β*), drawn randomly from the experimentally characterized distributions with mean *µ_δ or τ_* and standard deviation *σ_δ or τ_* (Figure 6b). These per-axon *α* and *β* values then serve as the means governing secondary, step-to-step variation. To capture instantaneous local fluctuations, the standard deviation at each step is derived from the average coefficient of variation (*ε* for step size and *ω* for turn rate), defined as the ratio *σ/µ* averaged across all empirical datasets.

At each time step, the instantaneous direction of the axon is drawn from a uniform distribution bounded by the experimentally characterized Katz dynamic ratios (Figure 5b-d). Once a kinematic case is stochastically selected, the trajectory advances by case-specific geometric rules (Figure 6c). For directed elongation (Case 1) and retraction (Case 3), movement is restricted to the longitudinal axis; the growth cone advances or retreats along its previous heading by the sampled local step size (*s*), with a turn rate (Δ*θ*) of zero. In contrast, during lateral wandering (Case 2), both the sampled local step size (*s*) and local turn rate (Δ*θ*) are applied simultaneously to compute the deflected coordinates of the next step. A summary of all empirical distributions and computational parameters utilized in these axonal outgrowth simulations is provided in Table 1.

**Table 1.** Summary of parameters used in the generative biased random walk simulation of axonal outgrowth. E/R/W represent elongation, retraction, and wandering ratios from Katz dynamics.

| Parameters | $\mu_\delta$ | $\sigma_\delta$ | $\mu_\tau$ | $\sigma_\tau$ | $\varepsilon$ | $\omega$ | Katz ratio (E/R/W) |
| --- | --- | --- | --- | --- | --- | --- | --- |
| PDL Phase 1 | 0.357 | 0.132 | 0.153 | 0.026 | 0.83 | 0.66 | 0.59/0.17/0.24 |
| PDL Phase 2 | 0.339 | 0.113 | 0.142 | 0.028 | 0.85 | 0.71 | 0.63/0.16/0.21 |
| PDL-LA Phase 1 | 0.350 | 0.131 | 0.152 | 0.032 | 0.84 | 0.64 | 0.60/0.15/0.26 |
| PDL-LA Phase 2 | 0.436 | 0.128 | 0.127 | 0.031 | 0.78 | 0.75 | 0.60/0.15/0.26 |

### 4.2 Simulating Substrate-Dependent Topology with Experimental Validation

To examine how substrate type shapes the topology of the axonal network during extended culture periods, we performed generative biased random walk simulations of axonal outgrowth at DIV 8, DIV 10, and DIV 13 for both PDL and PDL-LA substrates (Figure 7). The simulation timeline was aligned with the experiment by initializing axonal outgrowth at DIV 6.5, which corresponds to DIV 6 plating followed by a 12-hour somatic adhesion window during which no outgrowth beyond the neurosphere is assumed to occur. Phase 1 parameters governed the simulation from DIV 6.5 to DIV 8, after which the model switched to Phase 2 parameters to capture the mature growth regime. At an early time point (DIV 8), the simulated outgrowth was largely indistinguishable between the two substrates, consistent with the comparable Phase 1 kinematic parameters measured experimentally (1). However, by DIV 13, a clear substrate-dependent divergence emerged; PDL-LA produced more extensive axonal projections than PDL. This divergence cannot be attributed to a single parameter, but reflects the combined Phase 2 kinematics, in which PDL-LA exhibits a larger step size (*µ_δ_* = 0.436 vs. 0.339 *µm/min*) and a lower turn rate (*µ_τ_* = 0.127 vs. 0.142 *rad/min*), despite PDL showing a higher elongation ratio. Across both substrates, simulated axonal density expanded monotonically with culture duration, reproducing the progressive radial outgrowth observed in our experimental neurosphere cultures and confirming that substrate-induced differences in single-axon kinematics translate predictably into macroscopic network architecture.

**Fig. 7.**
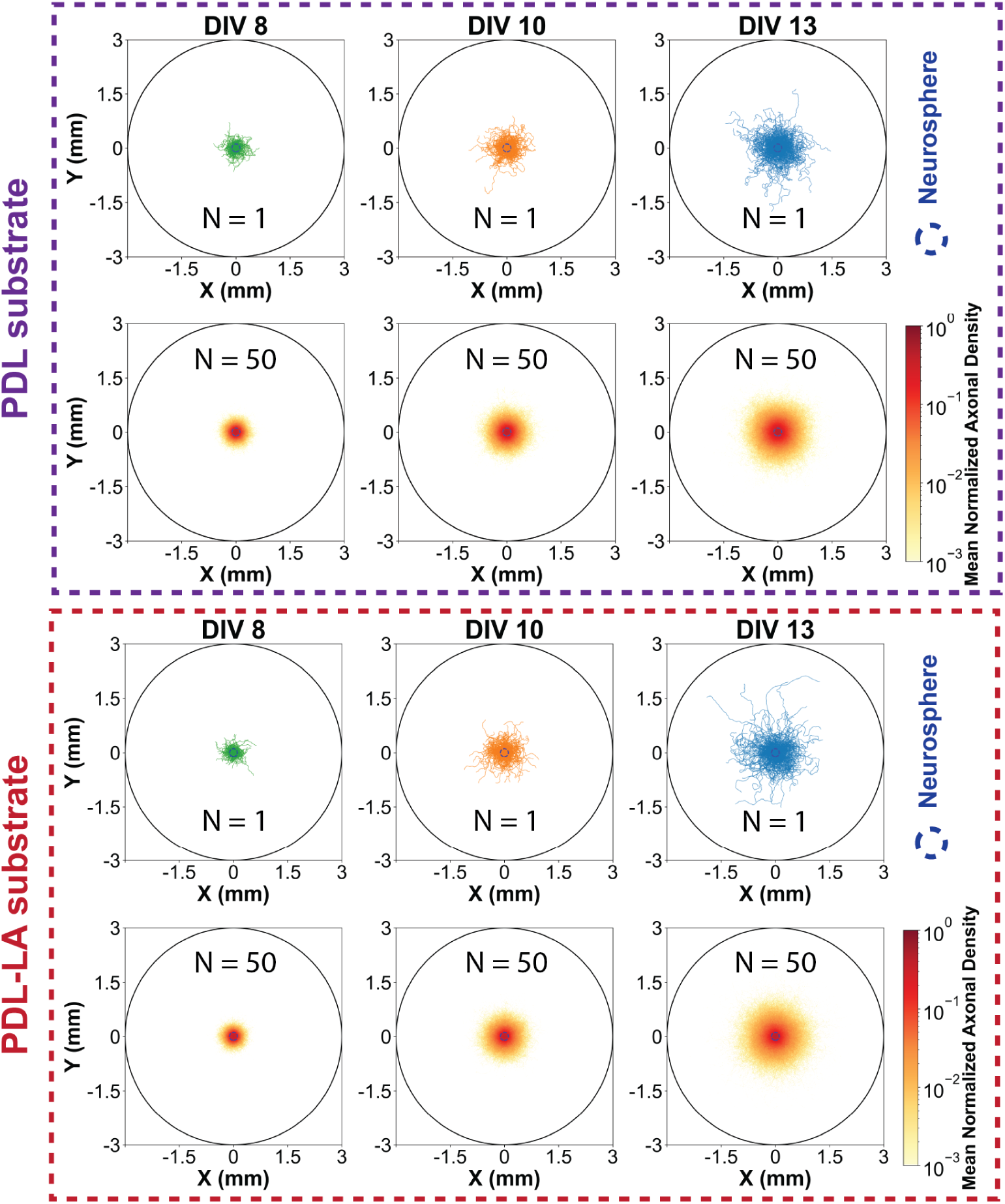
Generative biased random walk simulations of axonal outgrowth over extended culture periods. Simulated axonal trajectories and spatial densities are shown for DIV 8, 10, and 13. To synchronize the model with the experimental timeline, axonal outgrowth is initiated at DIV 6.5, accounting for plating at DIV 6 followed by a 12-hour somatic adhesion period, with no growth assumed prior to this checkpoint. Phase 1 parameters govern the simulation from DIV 6.5 to DIV 8, after which the model switches to Phase 2 parameters for the remainder of the culture period. In all panels, the central blue dotted boundary describes the initial axonal seeding coordinates. Each visualization overlays a single representative stochastic trajectory ensemble onto a spatial heatmap of the mean normalized axonal density, averaged across 50 independent simulations per experimental condition.

To quantitatively validate the generative model against experimental ground truth, we compared the simulation output to two independent empirical datasets that were not used in generating the distributions the model draws from at both macroscopic and microscopic scales (Figure 8). At the macroscopic scale, the normalized radial density profile of simulated axonal outgrowth at DIV 13 closely followed the normalized *βIII*−tubulin (TUBB3) fluorescence intensity profile measured from DIV 13 neurosphere cultures on PDL substrates (Figure 8a-b), confirming that the model reproduces the spatial extent and decay of axonal density observed in fixed cultures. At the microscopic scale, the trajectory coordinates (*x, y*) extracted from the simulation were used to recompute the underlying kinematic metrics and compared to the corresponding experimental distributions. Violin-plot comparisons revealed no statistically significant differences between the empirical and simulated populations for global step size, trajectory directionality, or the three Katz dynamic ratios (elongation, wandering, and retraction) (Figure 8c-g; n.s., *p >* 0.05). The simultaneous agreement at both scales demonstrates that the hierarchical, two-tiered random walk framework captures not only the statistical fingerprint of individual axonal kinematics but also the emergent macroscopic network architecture they produce, establishing the model as a quantitatively predictive tool for substrate-dependent axonal outgrowth.

**Fig. 8.**
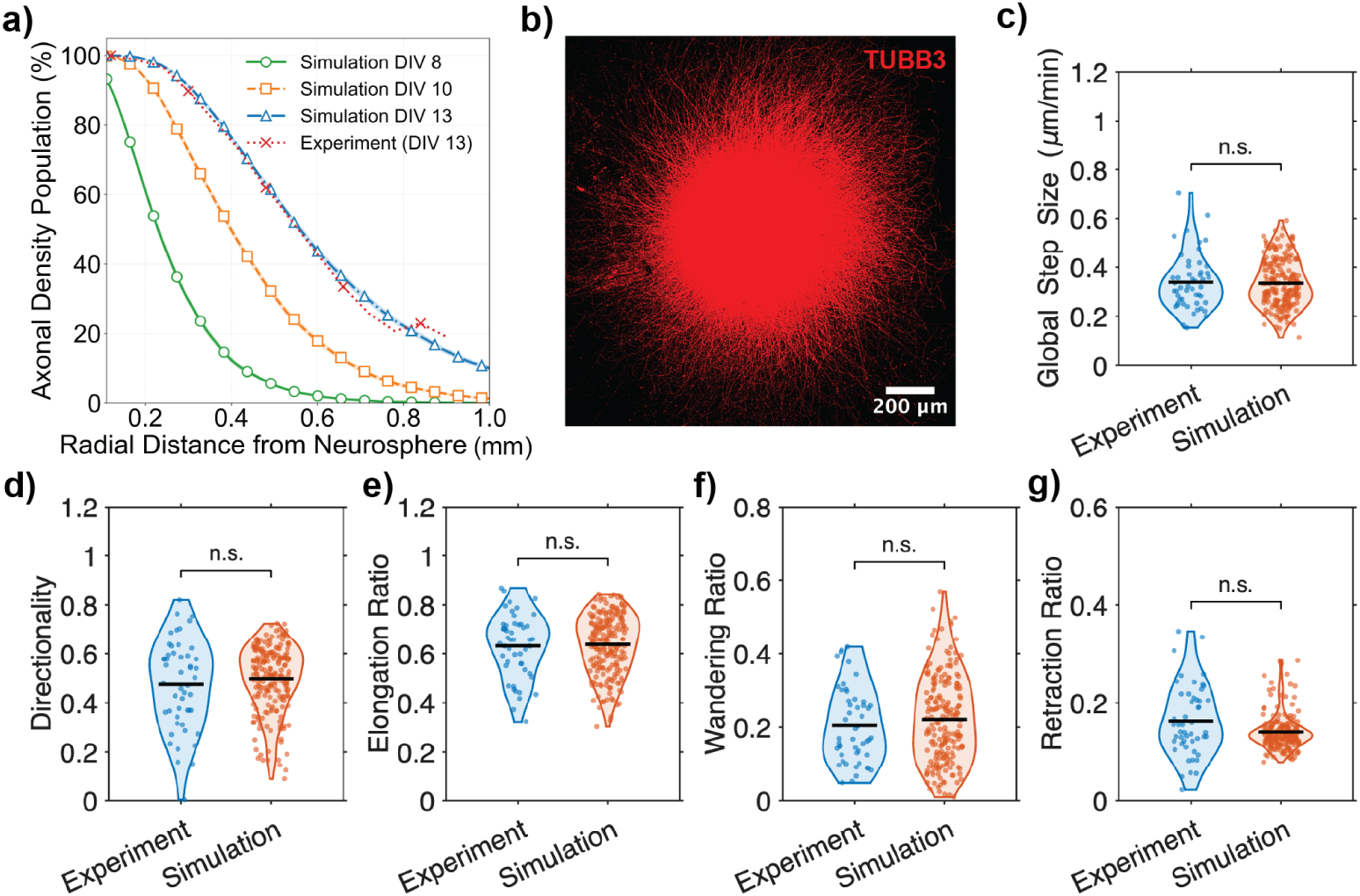
Macroscopic and microscopic experimental validation of the generative axonal growth simulation. a) Comparison of macroscopic radial axonal density profiles. Simulated axonal density populations are plotted for DIV 8 (green), DIV 10 (orange), and DIV 13 (blue); the red dashed line shows the normalized experimental fluorescence intensity profile from *βIII−*tubulin (TUBB3) staining of DIV 13 neurosphere cultures on PDL substrate. b) Representative immunocytochemistry (ICC) image of a DIV 13 neurosphere stained for TUBB3 (red), which served as the experimental ground truth for the normalized intensity profile mapped in a). c-g) Microscopic kinematic validation. Raw (x,y) trajectory coordinates generated by the simulation were analyzed to extract dynamic metrics for direct comparison with the experimental dataset. Violin plots confirm statistically no difference between the empirical and simulated populations for c) global step size, d) directionality, and the Katz dynamic ratios: e) elongation, f) wandering, and g) retraction (n.s., not statistically significant: *p >* 0.05).

## 5 Discussion

In this study, we established a semi-automated interactive tracking pipeline for quantifying neurite dynamics in mouse cortical neurospheres cultured on functionalized substrates, and used the resulting empirical kinematic distributions to construct and validate a generative biased random walk model of cortical neuron axonal outgrowth. The pipeline extracts both morphometric (step size, turn angle, directionality) and dynamic metrics based on a stochastic-state decomposition (elongation, lateral wandering, retraction), capturing axon kinematics at minute-scale temporal resolution. From these measurements, we constructed a two-tiered hierarchical model that captures both global axon-to-axon and local step-to-step variability, reproducing single-axon kinematics at the microscopic scale and neurosphere outgrowth topology at the macroscopic scale.

A key question for interpreting our results is the extent to which the kinematics of neurosphere-derived neurites generalizes to the broader cortical-neurite population. Most quantitative axonal-growth measurements in the literature [16, 58] derive from dissociated single-neuron cultures, in which isolated growth cones extend in a sparse, low-population density environment. In contrast, neurites emerging from a neurosphere are likely to initially traverse a dense intrasphere environment before extending radially into the surrounding substrate, and therefore the resulting kinematics may differ systematically from those of dissociated neurons. Indeed, Rizzo *et al.* reported axonal velocities of up to ∼ 0.21*µm/min* for rat cortical neurons on PDL-coated glass [58], and Smirnov et al. reported an average growth velocity of ∼ 0.2 *µm/min* for cerebellar granule neurons on laminin-coated substrates [16]. Our neurosphere-derived mean step sizes (0.34 − 0.44 *µm/min*) exceed these single-neuron values, and faster outgrowth has also been reported for larger spheroids (0.11 ± 0.07 *µm/s* in a 2.5 *mm i*^3^ neurosphere) [59]. This spheroid-associated acceleration may be attributed to two neurosphere-specific factors: 1) axon fasciculation, which provides mechanical guidance and reduces growth-cone reorientation costs through axon-axon adhesion mediated by L1-CAM and NCAM [60]; and 2) the more mature developmental window (DIV 6.5+) compared with the embryonic dissociated cultures used in the above studies. Despite this systematic offset in absolute speed, the shape of our step-size distribution remains consistent with the heavy-tailed, non-Gaussian distributions previously reported for cortical neurite-kinematics [58], suggesting that the underlying stochastic structure of growth-cone dynamics is preserved in the neurosphere context even when the mean speed is elevated.

A particularly interesting finding is the phase-specific substrate dependence; during Phase 1 (DIV 6.5-7.5), the kinematic parameters on PDL and PDL-LA are statistically indistinguishable, whereas during Phase 2 (DIV 8-9), PDL-LA neurites exhibit a significantly larger step size and a lower turn rate than PDL neurites (1). This biphasic substrate effect is consistent with the molecular-clutch mechanism of laminin-mediated mechanotransduction described by Abe et al. [52], in which the L1-CAM-laminin interaction provides a mechanical clutch that couples retrograde actin flow in the growth cone to the substrate, generating forward traction. In this framework, Phase 1 represents an adhesion-establishment period during which integrin and L1-CAM receptors are still being recruited to the growth-cone membrane. Until clutch engagement reaches a critical threshold, neurites on both substrates behave indistinguishably, dominated by PDL-mediated nonspecific electrostatic adhesion. Once laminin-specific clutches are fully engaged in Phase 2, the resulting traction increase produces faster forward advance and more directionally persistent locomotion, as stronger substrate coupling stabilizes growth-cone orientation against cytoskeletal fluctuations. This biophysical interpretation is consistent with prior findings that laminin-integrin engagement activates FAK at point contacts, which in turn modulates Rho-family GTPase activity (Rac1 and Cdc42) to drive membrane protrusion and stabilize growth-cone adhesion to the substrate [61, 62]. Under this signaling regime, growth cones can extend faster and reorient less frequently on uniform laminin substrates, because stable point contacts maintain a consistent traction axis [63].

The combination of our interactive tracking pipeline and the validated generative random walk model establishes a quantitative platform for investigating neurite outgrowth across a wide range of *in vitro* paradigms. To our knowledge, this is the first study to systematically characterize cortical neurosphere-derived neurite dynamics across functionalized substrates while tightly controlling other experimental variables: transfer day, neurosphere number, and neurosphere size, yielding highly reproducible results that provide a robust empirical ground truth for downstream applications. First, the framework can serve as the foundation for digital-twin models of neurodevelopmental [64] or neurodegenerative [65] diseases, in which neurospheres derived from patient-iPSC lines [59] modeling Alzheimer’s or Parkinson’s disease are characterized by refitting the model’s kinematic parameters to disease-specific imaging data, enabling rapid *in silico* screening of dose-response and time-course effects for candidate drugs and neurotrophic factors against a well-defined kinematic baseline. Second, the model can be used in a forward-design mode to optimize the geometry of neurosphere-based bioengineered circuits, predicting how inter-neurosphere spacing, reservoir size, and axon-penetration channel geometry jointly determine the timing and probability of structural connectivity between neurospheres before fabrication. This forward-prediction capability is particularly valuable for neuronal modular circuit networks [31–33], which are gaining attention in the emerging field of biological intelligence [38, 39, 66] as alternatives to conventional AI hardware platforms, and whose structure-function relationships remain incompletely understood. Together, the two applications above position our framework as a principled tool that unifies disease modeling and circuit engineering, advancing the *in vitro* neural system from descriptive observation to predictive, design-driven outcomes.

However, some limitations are worth noting. The current framework treats each axon as a single, non-branching trajectory, yet axonal branching is a biophysically and functionally fundamental phenomenon [67, 68]. Interstitial and terminal branching are the primary mechanisms by which a single axon tends to multiple branches and forms terminal arbors, allowing vertebrate neurons to integrate information from divergent regions of the nervous system [69, 70]. Branching could be incorporated into the random walk framework as a birth-death process, in which branch addition and elimination occur at rates estimated directly from time-lapse tracking data [71]; this extension would allow the model to capture true network connectivity rather than radial outgrowth alone. Complementary approaches such as phase field methods solved with isogeometric analysis capture the full spatial morphology of branching neurite tree directly, offering an alternative route to structural realism at higher computational cost [72, 73]. A second limitation is that our model assumes a spatially uniform environment, whereas real axonal growth reflects an interplay between deterministic biases (from chemical gradients, substrate mechanics, and geometry) and stochastic fluctuations (from receptor binding, signaling, adhesion, and cytoskeletal dynamics) [13, 74–76]. A Langevin and Fokker-Planck formulation could be a natural way to capture this interplay: the Langevin equation models each growth-cone trajectory as a deterministic drift term (directional bias from external cues) plus a stochastic noise term (intrinsic motility variability), while the equivalent Fokker-Planck equation describes how the probability density of the position of the axon evolves over time [77]. This continuous framework offers several advantages over discrete random walk models; it separates directional bias from motility noise, provides a mechanistically interpretable link between single-axon dynamics and population density, and enables semi-analytic prediction of axonal density in arbitrary cue fields and patterned substrates, avoiding the trajectory-by-trajectory Monte Carlo sampling our current model requires. Combining stochastic branching with a continuous biophysical formalism in future studies could yield a fully biophysics-based model capable of predicting how axonal networks self-organize under combined chemical, mechanical, and geometric constraints. Such a model would enable realistic *in silico* simulation of neurite growth and, in turn, support the rational design of diverse modular neural circuit architectures.

## 6 Conclusion

In this study, we have addressed a fundamental challenge in quantitative analysis of *in vitro* neurite outgrowth, namely the difficulty of disentangling intrinsic growth cone dynamics from confounding cell body motility in conventional dissociated 2D single-neuron cultures, by leveraging mouse cortical neurospheres as an experimental model with a fixed spatial origin, controllable size, and tightly constrained transfer day and seeding parameters. We have developed a semi-automated interactive neurite-tracking algorithm to extract neurite kinematic metrics across two developmental phases on two functionalized substrates (PDL and PDL-LA), revealing a notable phase-specific substrate divergence. Building on these measurements, we have constructed a generative biased random walk model with a two-tier hierarchical structure capturing both global axon-to-axon and local step-to-step variability, parameterized by experimentally measured step size, turn rate, and Katz dynamic ratio distributions. The model reproduces neurite kinematics at the microscopic scale and neurosphere outgrowth topology at the macroscopic scale, validated against fixed-timepoint TUBB3 immunocytochemistry. Together, this work integrates a controlled experimental model, a quantitative tracking pipeline, and a multi-scale generative simulation into a single coherent framework, advancing *in vitro* neurite analysis from descriptive characterization toward predictive, design-driven modeling. We anticipate broad applicability of this framework to disease-specific digital twins constructed from patient-iPSC neurospheres, and to the rational forward-design of multi-neurosphere modular circuits in which structural connectivity is specified at the design stage.

## Information Sharing Statement

The data set presented in this work and the code used to analyze the data are available at https://github.com/CMU-BORG/CBET Neurosphere Tracking-and-Analysis.git and archived at https://doi.org/10.5281/zenodo.21249734.

## Acknowledgments

The authors would like to acknowledge use of BioShare facilities (1f3e03) and its staff support. The manuscript and all analysis scripts were written by the authors. Generative AI was used solely to review and edit this author-generated content. All AI-suggested revisions were inspected and validated by the authors, and no AI tool was used to generate or alter data, figures, or results. The authors take full responsibility for the content of the publication.

## Author Contributions

Conceptualization: CK, VWW; Data Curation: (Lead) CK, (Supporting) MK, HC; Formal Analysis: CK; Acquisition of funding: VWW, YJZ; Investigation: CK; Methodology: CK, MK, VWW; Software: CK; Supervision: VWW, YJZ; Visualization: CK; Writing - Original Draft: CK; Writing - Review Editing: CK, MK, HC, TH, YJZ, TCK, VWW

## Funding

This project was supported by US National Science Foundation grant CBET-2332084 and Manufacturing PA Innovation Program. T. Y. Hsieh and Y. J. Zhang were also supported by the US National Science Foundation grant CMMI-1953323.

## Declarations

### Ethics Approval

Not Applicable

### Consent to Participate

Not Applicable

### Competing Interests

The authors have no competing interests to declare that are relevant to the content of this article.

## Notes

https://github.com/CMU-BORG/CBET_Neurosphere_Tracking-and-Analysis.git

https://doi.org/10.5281/zenodo.21249734

### Competing Interest Statement

The authors have declared no competing interest.

